# Neonatal White Matter Microstructure Predicts Infant Attention Disengagement from Fearful Faces

**DOI:** 10.1101/2025.05.28.655764

**Authors:** Hilyatushalihah K. Audah, Elmo P. Pulli, Saara Nolvi, Ashmeet Jolly, Aylin Rosberg, Silja Luotonen, Isabella L.C. Mariani Wigley, Niloofar Hashempour, Ru Li, Elena Vartiainen, Ilkka Suuronen, Tuomo Häikiö, Harri Merisaari, John D. Lewis, Riikka Korja, Jetro J. Tuulari, Hasse Karlsson, Linnea Karlsson, Eeva-Leena Kataja

## Abstract

Infants develop an attentional bias towards faces already at birth, with further specification towards fearful faces emerging at 6 months and diminishing around 11 months of age. However, the neurobiological origins of attentional bias to fear are still poorly understood. To understand the neural structures underlying perception of facial expressions, the current study utilized newborn diffusion magnetic resonance images (N = 86; 41 females; μ *=* 27.15 days) and eye tracking from the same infants at 8-months (μ = 8.75 months) as a behavioural measure. An overlap paradigm was used to measure attention disengagement from fearful, happy, and neutral faces. Tract-based spatial statistics revealed that higher white matter (WM) mean diffusivity in widespread regions across the brain was associated with lower attention disengagement from fearful faces. The same association was found with happy faces but was limited to only the splenium of the corpus callosum and sensorimotor pathways. Variance in neonatal WM microstructure may reflect individual differences in growth that is related to attentional bias development later in infancy.

---

Facial expressions serve as vital cues in social interactions facilitating communication (Pascalis & Kelly, 2009) and emotional understanding. Already at birth, infants prefer tracking face-like stimuli compared to non-face stimuli (Farroni et al., 2013; Johnson & Tucker, 1996; Valenza et al., 1996). Attention orienting —the process of directing attention towards specific stimuli— becomes functional 3 to 6 months after birth (Amso & Johnson, 2008; Simion et al., 2007). Complex attention orienting behaviour, including suppressing competing stimuli and disengaging attention begins to appear after 4 months (Amso & Scerif, 2015; Butcher et al., 2000). A preference for specific affective faces emerges in 6- month-old infants, who focus more on fearful faces than on happy or neutral faces (Kataja et al., 2022; Peltola et al., 2009). This attentional bias (AB) to fear in infants diminishes around 11 months (Leppänen, 2016; Peltola et al., 2013) possibly to facilitate semi-independent motoric exploration of the infant away from the caregiver (Callaghan et al., 2019).

Research suggests that neonatal neural structure can serve as a precursor to later behavioural phenotypes and cognition. In newborns, increased left amygdala volumes were related to heightened infant attention disengagement from fearful faces at 8 months (Tuulari et al., 2020). Stronger neonatal amygdala functional connectivity with the anterior cingulate and insula predicted higher fear and more advanced cognitive development in 6- month-old infants (Graham et al., 2016). In adults, the amygdala is well established as part of a multi-network collaboration, processing socio-emotional information (Meisner et al., 2022), central to threat detection and aversive learning (Costafreda et al., 2008; Kong & Zweifel, 2021) alongside the hippocampus and parts of the cingulate cortex (Palomero-Gallagher & Amunts, 2022).

Amygdala lesions are linked to various social and non-social deficits (Gallagher & Chiba, 1996; Pitteri et al., 2019). The local fibre density of a functionally afferent pulvinar-amygdala pathway significantly predicts fearful face recognition ability, providing evidence of a subcortical pathway related to fear recognition in adults (Mcfadyen et al., 2018). However, the neural pathways underlying socioemotional processes in early life remains unclear. More research is needed to explore the developmental origins of fear and its neural markers by studying infants. Specifically, diffusion tensor imaging (DTI) of white matter (WM) tracts can provide insight into the role of structural networks in facilitating emotional processing within typical development, facilitating the detection of aberrant behaviour and its underlying mechanisms.

The cingulum (CNG) is one potential pathway underlying attention towards facial expressions in early life. The CNG is a major tract involved in fear processing, connecting the cingulate gyrus with subcortical structures such as the amygdala, with the frontal, parietal and medial temporal cortices (Bubb et al., 2018). In adults, higher fractional anisotropy (FA) in the left CNG is related to increased posttraumatic stress disorder (PTSD) symptom severity (Averill et al., 2018), where maladaptive attentional orienting is a hallmark of the disorder (Schoorl et al., 2014).

In preterm neonates, right uncinate fasciculus (UF) microstructure predicted better emotion regulation abilities at 4 years of age (Kanel et al., 2021). The UF at 3-months, was the most robust predictor of negative emotionality regulation in 9-months infants compared to the cingulum and forceps (Banihashemi et al., 2020). In adults, left UF integrity has been associated with greater AB to briefly presented fearful faces (Carlson et al., 2013). Due to its apparent involvement in affect regulation, the UF may also influence facial fear processing in infants.

The inferior longitudinal fasciculus (ILF) is an alternative pathway for facial fear processing, connecting the occipital cortex with anterior temporal lobes, fusiform face area, and the amygdala (Herbet et al., 2018). In adults, faster emotion discrimination for fearful faces was associated with higher FA values in the ILF, but lower functional connectivity with the face processing network and the right amygdala (Marstaller et al., 2016). Damage to the ILF and the inferior fronto-occipital fasciculus (IFOF) can result in poorer performance on a facial emotion identification test (Genova et al., 2014). This research suggests that WM integrity of the ILF is involved in fearful face processing. However, all this research was conducted in adults, and none in infants. Notably, the involvement of the ILF in visual-emotion processing may be reduced in infants due to the ILF’s late developmental window extending into adulthood (Dubois et al., 2008; Lebel & Beaulieu, 2011). More research is needed to further understand the ILF’s potential role in early life affective facial processing.

Additionally, another study found that lower FA values of the left inferior stria terminalis significantly predicted heightened fear at 6 months and higher subsequent increases in fear at 12 and 18 months (Planalp et al., 2022). Limbic structures such as the fornix and stria terminalis are the earliest tracts to develop in neonates (Yap et al., 2013) and hence may also influence fear processing in infants. However, the Infant Revised Behaviour Questionnaire (Gartstein & Rothbart, 2003) used in the Planalp et al (2022) study contains items regarding infant behaviour towards novelty. Moreover, the infants’ behaviour was also reported by parents, whereas this study uses direct measures of infant behaviour through eye tracking. Hence, to our knowledge, there are no studies directly examining the neural structural pathways underlying facial expression perception processing from infant behaviour.

The current study aims to extend previous findings by focusing on neonatal structural pathways associated with infant attention towards facial expressions. This study will first use a region-of-interest (ROI) *a priori* approach to determine whether neonatal CNG, UF, and ILF are related to AB towards faces in 8-month-old infants. Subsequently, this study explored and identified the potential white matter structures involved in facial perception using a whole brain voxelwise approach extending the pre-registered research plan. The hypothesis was that the CNG, UF, and ILF can predict variance in infant fear bias, and that higher fear bias is associated with higher FA in these tracts.

## Methods

Data used in this study were collected in accordance with the Declaration of Helsinki. All studies were approved by the Ethics Committee of the Hospital District of Southwest Finland. Mothers gave informed consent on behalf of their infants. The pre-registered plan is available on the Open Science Framework (https://osf.io/exjr2).

### Participants

All data collected are part of the ongoing longitudinal FinnBrain Birth Cohort study (Karlsson et al., 2018). Pregnant mothers were recruited during their first ultrasound visit (12 gestational weeks; gwk) from three maternal welfare clinics in the Southwest Finland Hospital District and the Åland Islands. The inclusion criteria were an ultrasound verified pregnancy and sufficient knowledge of one of Finland’s official languages (Finnish or Swedish). At 2 to 7 weeks postpartum, a subsample of the whole cohort families was invited for magnetic resonance imaging (MRI) visits to the *FinnBrain Neuroimaging Lab.* At 8 months (corrected age from the due date) families participated in a larger neuropsychological visit to the *FinnBrain Child Development and Parental Functioning Lab*. A total of 189 infant-mother dyads participated, of these children, 96 infants had both newborn diffusion MRI and 8-months eye-tracking data. After quality control (QC) the total sample size was 86 participants (41 female). All scanned infants were at least 2500 gr at birth, born gwk 36 or later, and were 291-320 days old (μ = 305.71, SD = 7.24 days) on the day of scanning from conception. At the eye-tracking experiment, infants were 7.46 to 9.71 months old (μ = 8.74 months, SD = 0.39). A summary of the sample characteristics can be found in Table 1.

**Table 1.**
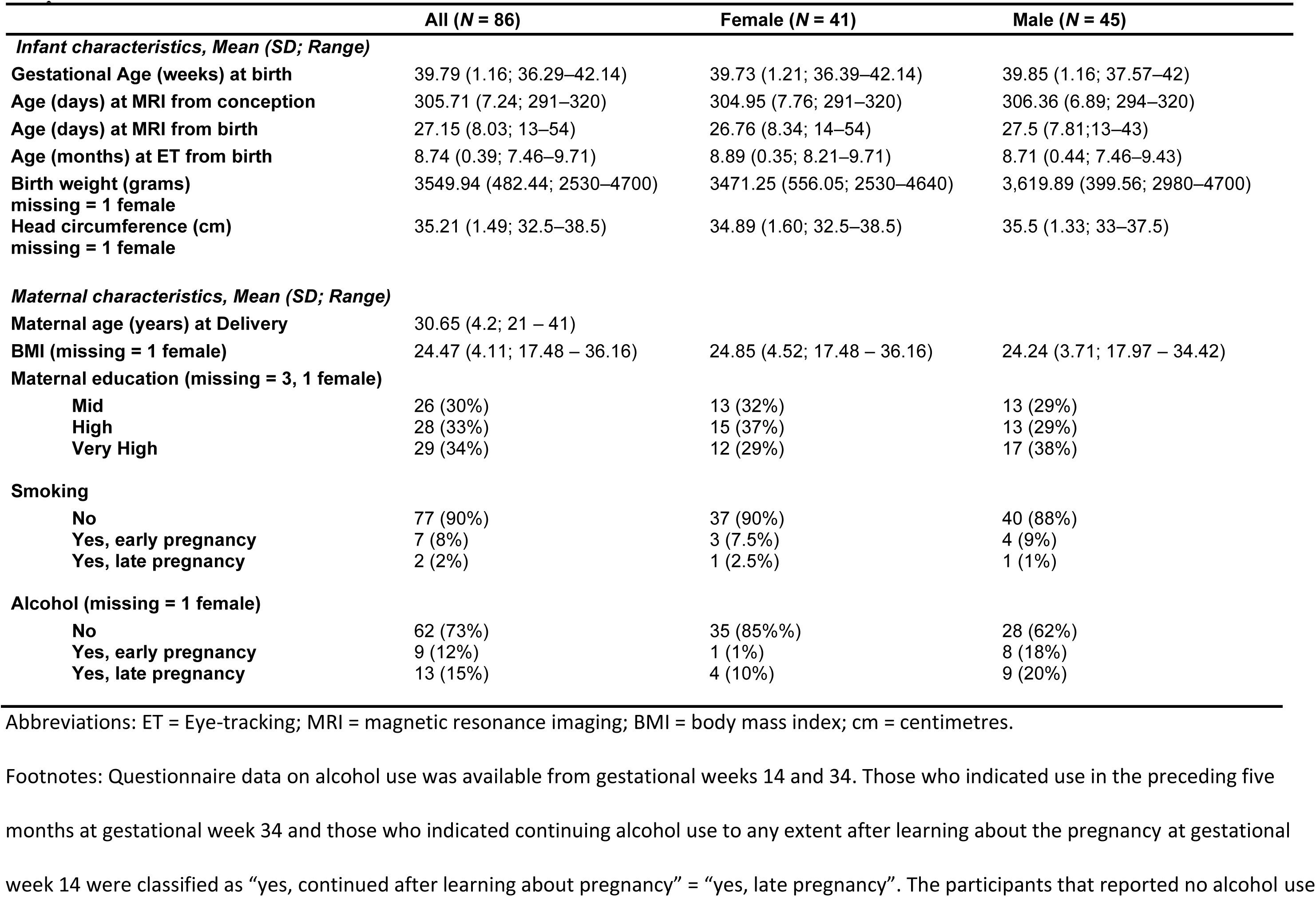
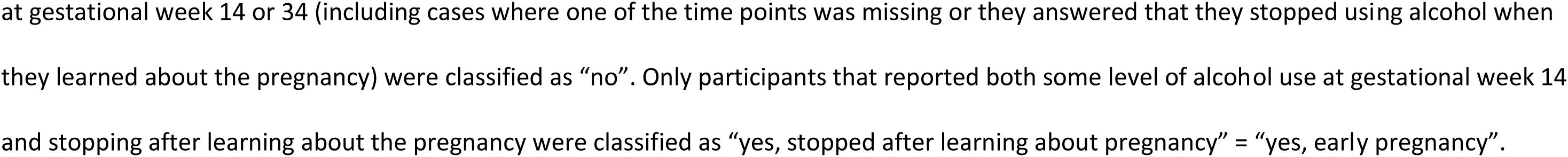
Sample characteristics.

|  | All (N = 86) | Female (N = 41) | Male (N = 45) |
| --- | --- | --- | --- |
| ***Infant characteristics, Mean (SD; Range)*** |  |  |  |
| **Gestational Age (weeks) at birth** | 39.79 (1.16; 36.29–42.14) | 39.73 (1.21; 36.39–42.14) | 39.85 (1.16; 37.57–42) |
| **Age (days) at MRI from conception** | 305.71 (7.24; 291–320) | 304.95 (7.76; 291–320) | 306.36 (6.89; 294–320) |
| **Age (days) at MRI from birth** | 27.15 (8.03; 13–54) | 26.76 (8.34; 14–54) | 27.5 (7.81; 13–43) |
| **Age (months) at ET from birth** | 8.74 (0.39; 7.46–9.71) | 8.89 (0.35; 8.21–9.71) | 8.71 (0.44; 7.46–9.43) |
| **Birth weight (grams)**<br>missing = 1 female | 3549.94 (482.44; 2530–4700) | 3471.25 (556.05; 2530–4640) | 3,619.89 (399.56; 2980–4700) |
| **Head circumference (cm)**<br>missing = 1 female | 35.21 (1.49; 32.5–38.5) | 34.89 (1.60; 32.5–38.5) | 35.5 (1.33; 33–37.5) |
| ***Maternal characteristics, Mean (SD; Range)*** |  |  |  |
| **Maternal age (years) at Delivery** | 30.65 (4.2; 21 – 41) |  |  |
| **BMI (missing = 1 female)** | 24.47 (4.11; 17.48 – 36.16) | 24.85 (4.52; 17.48 – 36.16) | 24.24 (3.71; 17.97 – 34.42) |
| **Maternal education (missing = 3, 1 female)** |  |  |  |
| Mid | 26 (30%) | 13 (32%) | 13 (29%) |
| High | 28 (33%) | 15 (37%) | 13 (29%) |
| Very High | 29 (34%) | 12 (29%) | 17 (38%) |
| **Smoking** |  |  |  |
| No | 77 (90%) | 37 (90%) | 40 (88%) |
| Yes, early pregnancy | 7 (8%) | 3 (7.5%) | 4 (9%) |
| Yes, late pregnancy | 2 (2%) | 1 (2.5%) | 1 (1%) |
| **Alcohol (missing = 1 female)** |  |  |  |
| No | 62 (73%) | 35 (85%%) | 28 (62%) |
| Yes, early pregnancy | 9 (12%) | 1 (1%) | 8 (18%) |
| Yes, late pregnancy | 13 (15%) | 4 (10%) | 9 (20%) |

Abbreviations: ET = Eye-tracking; MRI = magnetic resonance imaging; BMI = body mass index; cm = centimetres.
Footnotes: Questionnaire data on alcohol use was available from gestational weeks 14 and 34. Those who indicated use in the preceding five months at gestational week 34 and those who indicated continuing alcohol use to any extent after learning about the pregnancy at gestational week 14 were classified as “yes, continued after learning about pregnancy” = “yes, late pregnancy”. The participants that reported no alcohol use
at gestational week 14 or 34 (including cases where one of the time points was missing or they answered that they stopped using alcohol when they learned about the pregnancy) were classified as “no”. Only participants that reported both some level of alcohol use at gestational week 14 and stopping after learning about the pregnancy were classified as “yes, stopped after learning about pregnancy” = “yes, early pregnancy”.

Data on infant sex, birthweight, head circumference at birth, date of birth (used to calculate infant age at scan), maternal pre-pregnancy BMI, maternal age at delivery, and maternal smoking during pregnancy were acquired from the wellbeing services county of Southwest Finland (VARHA) records. Other background information was obtained from self-report questionnaires filled in by the mothers at gwk 14 and 34, including maternal alcohol consumption during pregnancy. Maternal education as a measure of socio-economic status (SES) was collected at gwk 14, with “middle education” defined as education up to upper secondary/vocational school, “high education” defined as education up to an applied University degree, and “very high education” defined as academic university degrees up to a doctorate/licentiate.

### MRI Acquisition

Infants were scanned using a Siemens Magnetom (Kumpulainen et al., 2020 for a more detailed description). Diffusion data were acquired using a standard twice-refocused Spin Echo–Echo Planar Imaging (SE-EPI) sequence, with a b-value of 1000s/mm^2^ and 2 × 2 × 2 mm^3^ isotropic resolution with a Field of View (FOV) of 208 mm. Data from infants with at least 20 diffusion-encoded directions were included in this study (Merisaari et al., 2023). A more detailed description of the MRI scanning protocol is available in previous reports (Lehtola et al., 2019; Merisaari et al., 2019, 2023; Tuulari et al., 2024).

### DTI analysis

MRI data preprocessing procedures are outlined in previous work (Kumpulainen et al., 2023; Merisaari et al., 2019, 2023; Rosberg et al., 2025). ROI-based analysis analyses were part of the pre-registered plan. ROI-based analysis utilized the AutoPtx pipeline (De Groot et al., 2013) to first mask the DTI tensor images. Then mean values for each delineated tract were calculated from the masks for each participant. This resulted in the brain-derived metrics tracts masks. The processed data were then visually inspected to ensure that all tracts were present, correctly placed, and did not extend to cortical and subcortical grey matter. Out of 169 participants, 152 successfully passed the manual QC, of which 76 had valid eye tracking data. The remaining 17 were excluded due to unsuccessful AutoPtx processing. The pipeline resulted in absolute values of FA, mean diffusivity (MD), and radial diffusivity (RD) for each participant. Only the FA and MD values of the predetermined tracts of interest: the CNG, UF and ILF were used as the brain-derived measures in the statistical analysis.

The following description of whole-brain analysis was not included in the pre-registered plan. Whole-brain analysis was done using TBSS (Tract-Based Spatial Statistics, (Smith et al., 2006) from the FMRIB Software Library (FSL; Smith et al., 2004) by creating a skeleton mask from the full dataset (N = 169) to maximize signal-to-noise ratio. Firstly, diffusion images were brain-extracted using the Brain Extraction Tool (BET; Smith et al., 2004). A tensor model was then fitted to the raw diffusion data using DTIFIT from FDT (FMRIB Diffusion Toolbox). All subjects’ FA data were aligned into a study specific template created from the whole dataset (Merisaari et al., 2019, 2023) using the nonlinear registration tool FNIRT (FMRIB’s Nonlinear Registration Tool; Andersson et al., 2007a, 2007b) which uses a b-spline representation of the registration warp field (Rueckert et al., 1999). Next, the mean FA image was created and thinned to create a mean FA skeleton which represents the centres of all tracts common to the group. Each subject’s aligned FA and MD maps were then projected onto this skeleton and the resulting data were fed into voxelwise cross-subject statistics. Additionally, the neonatal JHU ICBM White Matter Labels Atlas (Mori et al., 2008) were transformed into the study-specific template space, binary masks were created and superimposed on the spatially normalized skeletonized data to identify WM tracts within the TBSS skeleton.

### Eye-tracking Procedure

The eye-tracking experiment took place in a dimly lit room. Infants sat on their parent’s lap approximately 50-70 cm away from the eye-tracker (EyeLink1000+, SR Research Ltd, Toronto, Ontario, Canada) and a computer display. The researcher sat in the same room but was separated from the infant and parent by a curtain during the assessment. The measurement was managed by the researcher using an independent computer. Before every measurement, a five-point calibration procedure was performed using audio-visual animation presented at 5 locations on the screen. Data on infants’ eye position was collected with a 500 Hz sampling frequency as infants viewed pictures of faces and non-faces on a computer screen.

### Paradigm

The overlap paradigm was used to study infant disengagement from a central stimulus to a lateral stimulus distractor (Figure 1). Stimuli included happy (HA), neutral (NE), and fearful (FE) facial expressions of two different women, with a scrambled face picture as the control stimulus (CS). One of these 4 stimuli was presented at the centre of the screen for 1000 ms. The lateral distractors were either a black and white checkerboard or circles, appearing for 3000 ms on the left or right side of the central stimulus (13.6° visual angle). One trial lasted for 4000 ms. The sizes of the central pictures were 15.4° × 10.8° and the distractor stimuli were 15.4° × 4.3°. After each trial, a visual animation was briefly shown to capture the infant’s attention back to the centre of the screen. Overall, the infants were shown 48 sets of trials, 12 trials per condition (each emotion type and a control picture), made up of 18 photographs of each woman and 12 CS, in a semi-random order.

**Figure 1.**
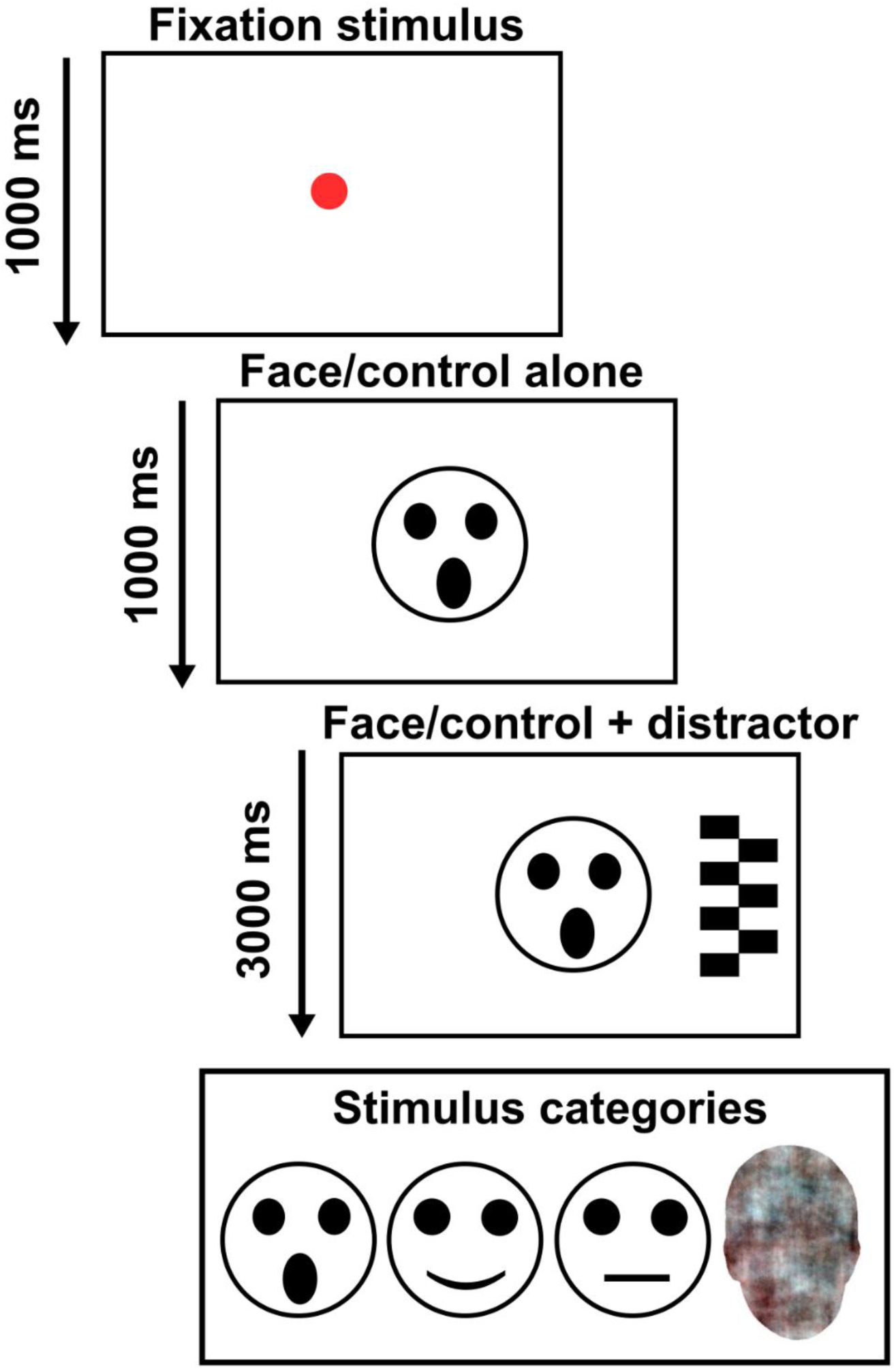
Illustration of overlap paradigm used in eye tracking. The task originally uses female faces, which has been anonymized for preprint.

### Eye-tracking data analysis

Eye-tracking data was analysed following the protocol described in a previous study (Leppänen et al., 2015) using MATLAB 2016b (The MathWorks Inc, 2016) functions (https://github.com/infant-cognition-turku/gazeanalysislib). The presence of a gaze shift from the centre to the lateral stimulus was determined using the XY coordinates of the infant’s gaze position. Infant eye tracking is often subjected to gaps (missing gaze points) and positional imprecision. QC criteria are:

i. the trial was excluded if the infant’s gaze was not in the area of the central stimulus for 70% of the time period that preceded possible disengagement from the central to lateral stimulus or if the infant gaze was not in the area at the end of the trial analysis period (1000 ms after lateral stimulus onset).
ii. trials with consecutive missing samples lasting 200 ms or longer were excluded.
iii. trials that were missing information on the exact timepoint of when the disengagement from the central stimulus to the lateral stimulus occurred were also excluded (i.e., the eye movement occurred during a period of missing gaze data).

The eye tracking analysis only included participants that had ≥ 3 valid trials for each stimulus condition. From the valid trials, the mean disengagement probabilities (DP) were calculated for each stimulus condition and used in the statistical analysis. The mean DP of non-fear faces (NF) was calculated using the average disengagement probabilities from neutral and happy stimuli (Mean NE+HA).

### Statistical analysis

Statistical analyses were conducted using IBM SPSS Statistics version 29 (IBM Corp., 2020) and JASP version 0.16.40 and version 0.19.1 (Love et al., 2019). Paired-samples t-tests were used to determine the presence of a face and fear bias in the whole sample. As per the pre-registered plan, Ordinary Least Square (OLS) regression models were conducted for each DP and bias scores with the CNG, UF, and ILF FA and MD values from the ROIs calculated with the AutoPtx pipeline separately for each hemisphere after confirming assumptions (normality of distribution, variance inflation factor, residual Q-Q plots). Covariates in the adjusted models differ slightly than in the pre-registered plan. The models are *FA or MD* ∼ *sex + gestational age (GA; weeks) + age at scan from conception (days) + age at eye tracking from birth (days) + maternal pre-pregnancy BMI + DP*. Pearson correlations were also conducted for all eye tracking DP conditions and confounding variables. All significance testing was 2-tailed with α = .05. A sensitivity analysis was performed with slightly different covariates than in the pre-registered plan.

Extending the pre-registered plan, whole brain voxelwise statistics of the FA and MD maps were conducted for all participants using FSL’s Randomise (Winkler et al., 2014) with 5000 permutations and threshold-free cluster enhancement (TFCE) corrected p-values. When creating the design matrix for the voxelwise statistics, any missing values (see Table 1 for details) were imputed with the sample mean for ratio variables and sample mode for categorical variables. FA and MD maps were analysed separately for face bias, fear bias, and for each condition (NE, FE, HA, CS). Additional whole brain voxelwise sensitivity analyses were conducted after the main analyses, further controlling for infant birthweight, head circumference, maternal age, education, alcohol and smoking during pregnancy.

Furthermore, a Sex × DP interaction analysis was done. Sex-based stratification was done by dividing the diffusion scalar data based on participant sex and then running whole-brain voxelwise statistics using the same covariates as the main analyses (results in Supplementary Materials).

## Results

### Whole brain voxelwise statistics

Whole-brain voxel wise statistics of the FA maps for each DP condition and bias scores did not show any significant results (p > .05). No significant results were found for fear bias scores, face bias scores, DP CS and DP NE conditions in the MD maps (p > .05). Higher MD was significantly associated with lower probability of disengaging from fearful faces (Figure 2, p ≤ .05) throughout the brain, but not occipital areas. These areas included the bilateral internal and external capsule, genu and body of the CC, bilateral superior fronto-occipital fasciculus (SFOF), bilateral anterior, superior, posterior corona radiata, left uncinate fasciculus, right superior longitudinal fasciculus (SLF). Higher MD was also significantly associated with lower probability of disengaging attention from happy faces (Figure 3, p ≤ 0.05) in the splenium of the CC. The Sex × DP interaction analysis did not yield any significant results.

**Figure 2.**
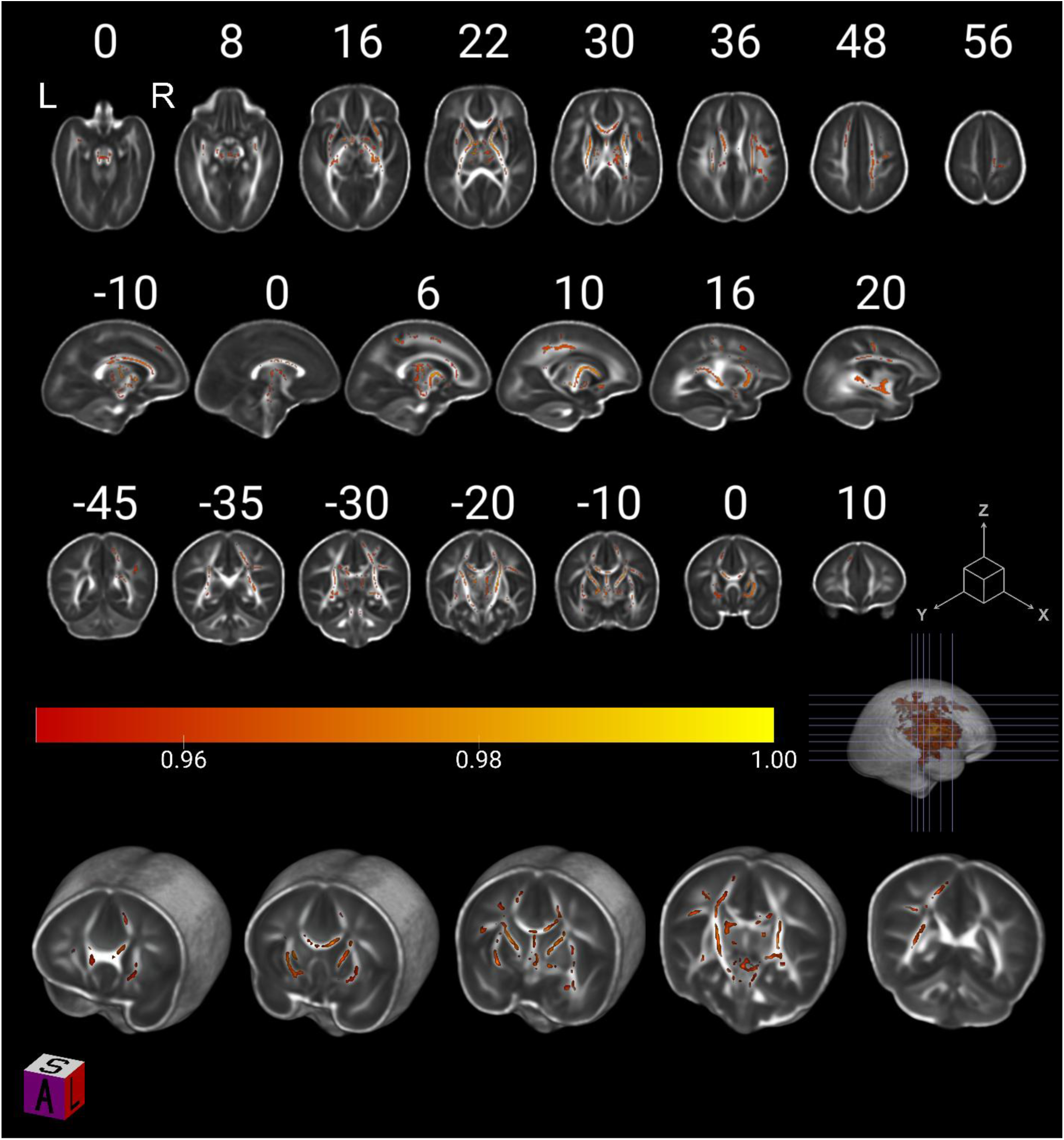
Higher mean diffusivity in newborn brains was associated with lower attention disengagement from fearful faces at 8 months (threshold-free cluster enhancement (TFCE) corrected p ≤ 0.05; across 5000 permutations). The colour bar represents 1 − *p*.

**Figure 3.**
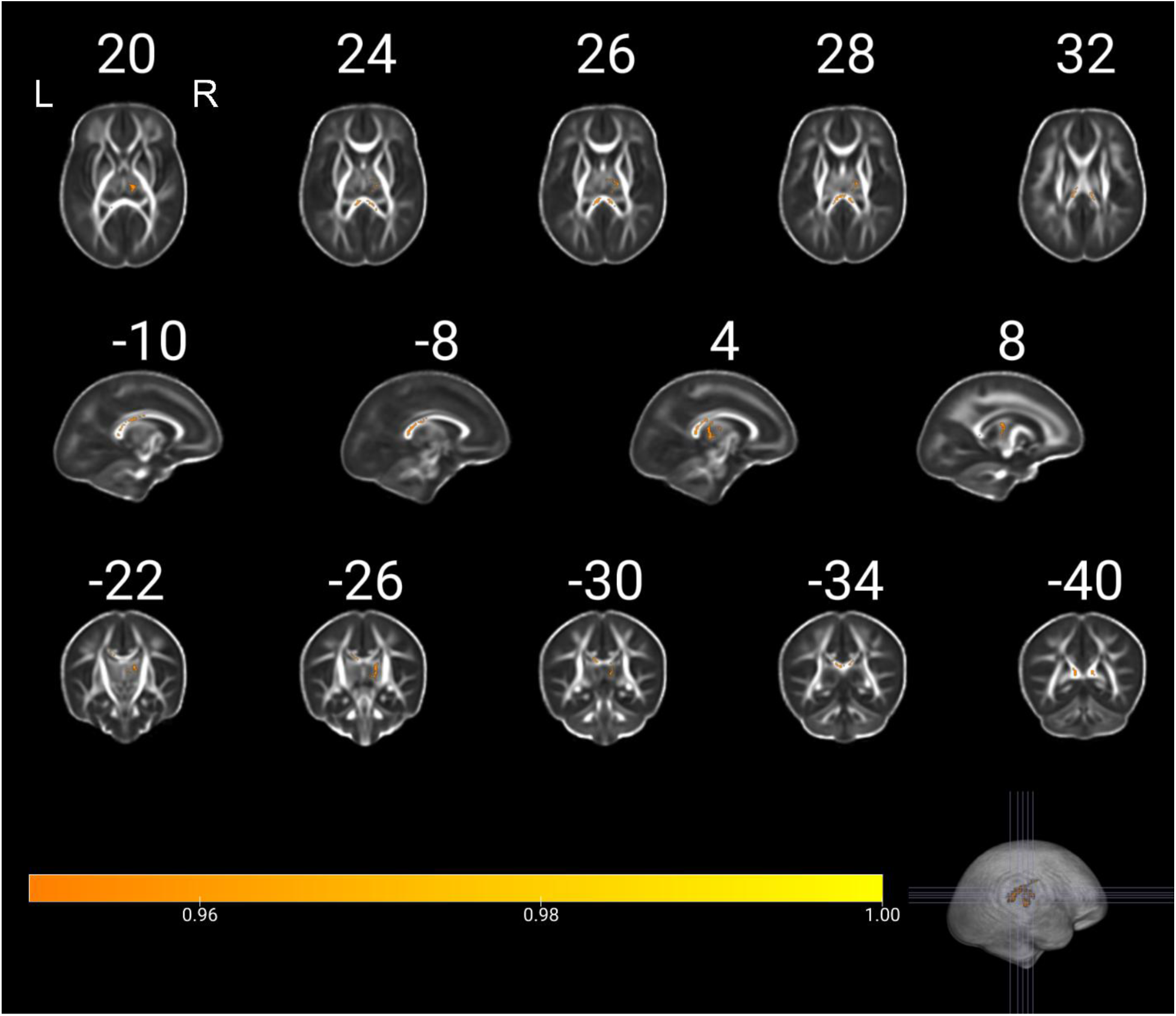
Higher mean diffusivity in newborn brains is associated with lower attention disengagement from happy faces at 8 months (threshold-free cluster enhancement (TFCE) corrected p ≤ 0.05; across 5000 permutations). The colour bar represents 1 − *p*.

### Attentional bias and ROI-based analysis

Paired samples t-tests showed a significant AB towards faces, t (85) = 7.684, p < .001, d = 0.131, when comparing DP CS (μ = .785) with the combined non-fear faces (μ = .792) consisting of DP NE (μ = .592) and DP HA (μ = .583). Infants were also significantly less likely to disengage attention from fearful faces (μ = .447) compared to non-fear faces, t (85) = −5.885, p < .001, d = 0.102. Pearson correlations (Table 2) showed that GA was significantly correlated with age at MRI (r = .460, p < .001) and age at ET (r = −.800, p <. 001) but not with other variables. Age at scan was significantly correlated with age at ET (r = −.300 p = .005). Maternal BMI was significantly negatively correlated with DP FE (r =−.230, p = 0.032). DP CS was significantly related to DP NE (r = .350, p = .001) and DP HA (r = .480 p < .001) but there were no other associations between all 4 DPs with infant characteristics. ROI-based analysis showed that the CNG, UF, and ILF did not significantly predict fear bias, face bias, or any of the DP conditions (Table S1.1). Sensitivity analyses did not yield any significant results.

**Table 2.**
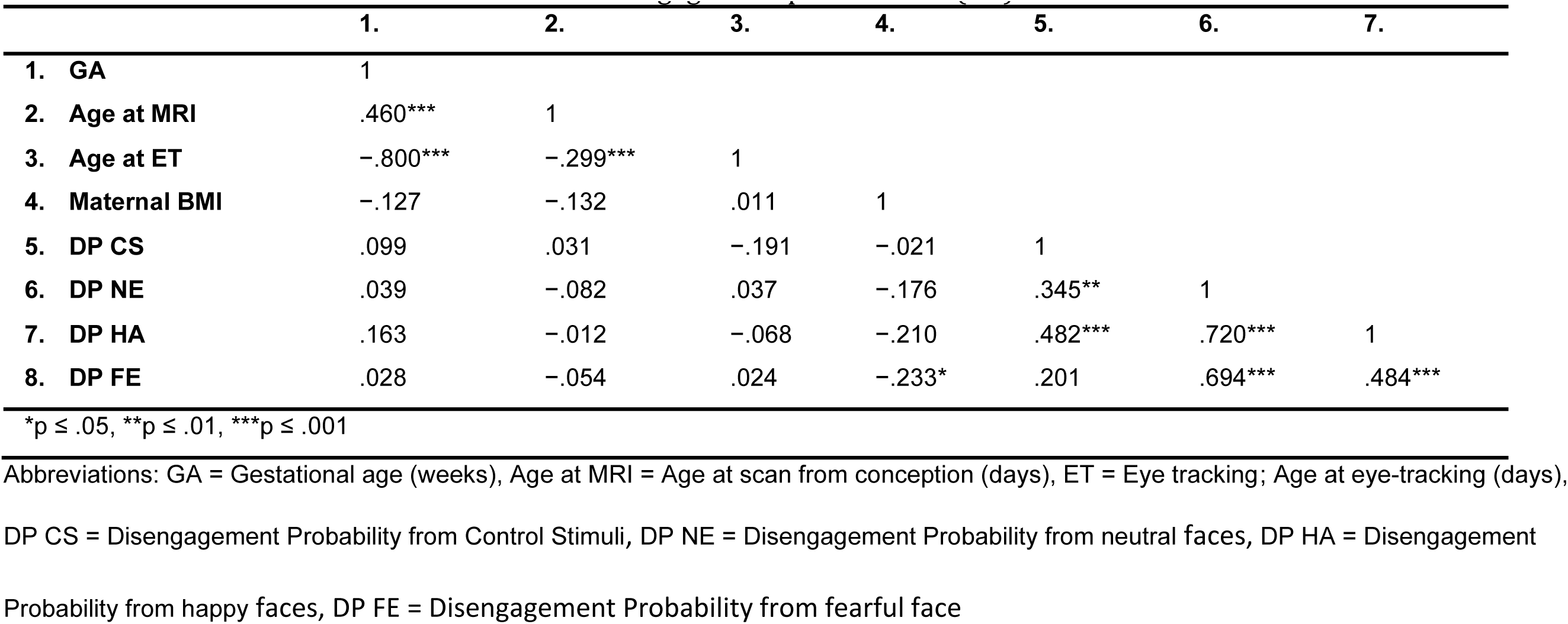
Pearson correlation matrix between attention disengagement probabilities (DP) and control variables.

|  | 1. | 2. | 3. | 4. | 5. | 6. | 7. |
| --- | --- | --- | --- | --- | --- | --- | --- |
| 1. GA | 1 |  |  |  |  |  |  |
| 2. Age at MRI | .460*** | 1 |  |  |  |  |  |
| 3. Age at ET | -.800*** | -.299*** | 1 |  |  |  |  |
| 4. Maternal BMI | -.127 | -.132 | .011 | 1 |  |  |  |
| 5. DP CS | .099 | .031 | -.191 | -.021 | 1 |  |  |
| 6. DP NE | .039 | -.082 | .037 | -.176 | .345** | 1 |  |
| 7. DP HA | .163 | -.012 | -.068 | -.210 | .482*** | .720*** | 1 |
| 8. DP FE | .028 | -.054 | .024 | -.233* | .201 | .694*** | .484*** |

\* $p \leq .05$ , \*\* $p \leq .01$ , \*\*\* $p \leq .001$
Abbreviations: GA = Gestational age (weeks), Age at MRI = Age at scan from conception (days), ET = Eye tracking; Age at eye-tracking (days),
DP CS = Disengagement Probability from Control Stimuli, DP NE = Disengagement Probability from neutral faces, DP HA = Disengagement
Probability from happy faces, DP FE = Disengagement Probability from fearful face

Figures 4 and 5 visualize the negative association, grouped by sex, between DP HA and DP FE with MD from the whole brain voxelwise statistics results. Sensitivity analyses changed some of the white matter areas implicated in the results. When including infant birthweight as a covariate of no interest into the DP FE voxelwise models more associations were found including in the occipital lobe. When controlling for head circumference, more associations were found in the splenium of the CC and in the right SLF, while maternal age showed less associations in inferior regions. With maternal alcohol consumption as a covariate of no interest we found more associations in the splenium of the CC. For DP HA, more associations were found in the posterior limb of the left internal capsule when controlling for birthweight or maternal age or alcohol consumption. All other covariates showed similar results to the main analyses or none. See supplementary materials for visualization.

**Figure 4.**
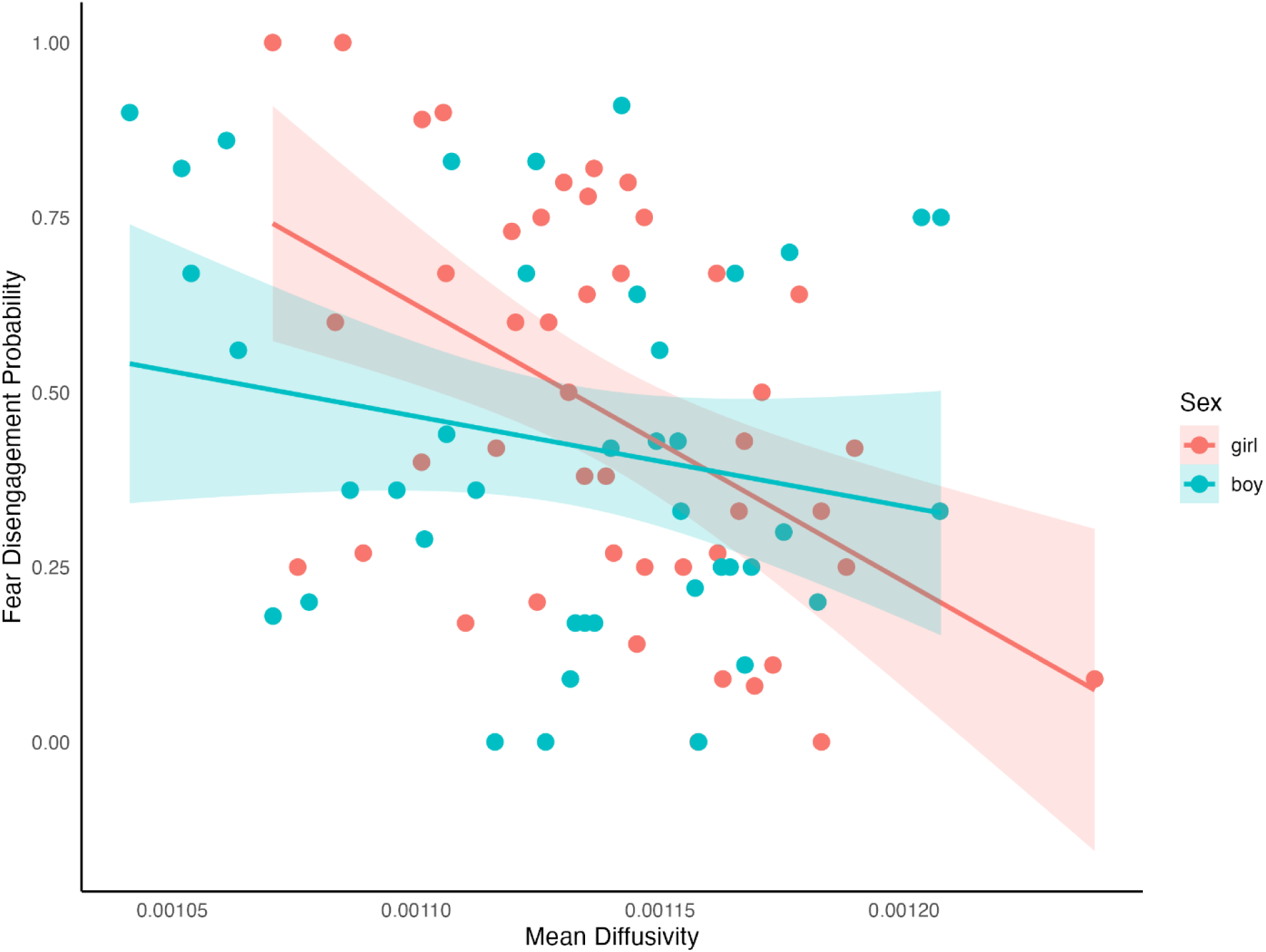
Scatterplot visualizing the negative association (p ≤ 0.05) between WM mean diffusivity across the whole brain and attention disengagement from fearful faces. These data are used for visualization purposes only, and the statistical inferences are based on FSL randomise whole brain voxelwise statistics.

**Figure 5.**
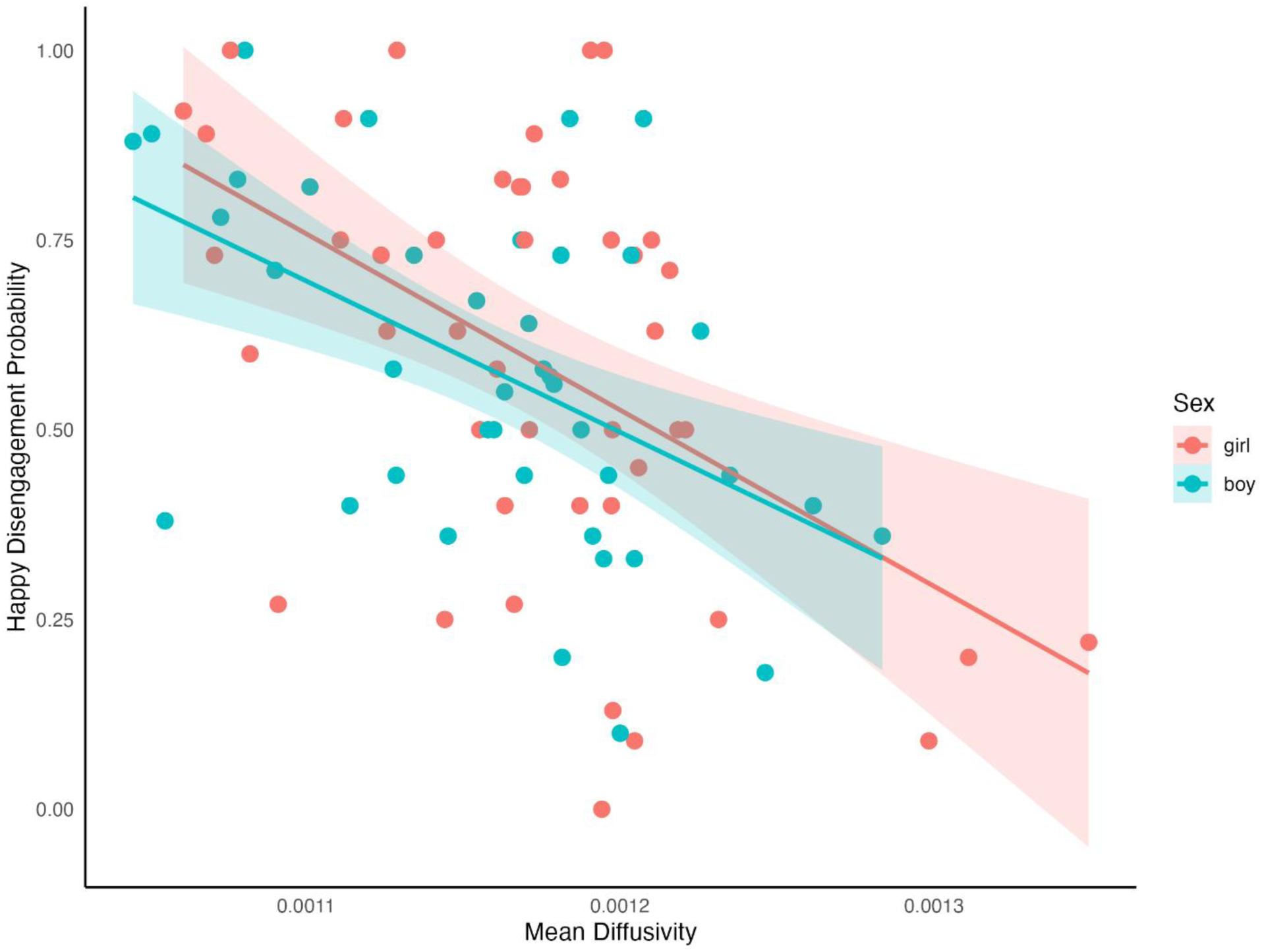
Scatterplot visualizing the negative association (p ≤ 0.05) between WM mean diffusivity across the whole brain and attention disengagement from happy faces. These data are used for visualization purposes only, and the statistical inferences are based on FSL randomise whole brain voxelwise statistics.

## Discussion

This study used diffusion-weighted MRI to identify neonatal WM tracts associated with an AB towards fearful faces 8-months old infants. The whole brain voxel-wise analysis showed that higher WM mean diffusivity in newborns was associated with sustained attention to fearful and happy faces during infancy. With fearful faces, higher MD was widespread throughout the brain, except in the posterior areas such as the occipital lobe. Meanwhile happy faces showed higher MD in the splenium of the CC. To our knowledge, this is a novel finding and provides further insight into how neonatal variance in WM microstructure is related to attentional biases in early life—behavioural phenotypes known to be relevant for socio-emotional development (Eskola et al., 2023; Peltola et al., 2018).

The whole brain voxelwise results extends on previous research examining infant attention that found higher FA underlies increased attention orienting behaviour in infants (Dowe et al., 2020). However, that study used hand puppets with large eyes and jingle bells as the attention-grabbing stimuli, which can be perceived as face-like stimuli, but also includes additional auditory stimuli (i.e., the bells)(Dowe et al., 2020) that may contribute to the findings differences in our study. Notably, FA and MD are different measures, and MD does not measure WM integrity. MD consists of the average radial, axial and another perpendicular component diffusivity; hence the increased diffusivity could indicate less myelination, less densely packed axons (Dubois et al., 2014), and smaller axonal radii (Pizzolato et al., 2023).

The association of heightened AB to facial expression and widespread MD found in this study is expected in rapidly developing infants. The findings align with the understanding that infant behaviours are closely linked to their overall developmental stage which is also reflected in neurodevelopment (Zhuo et al., 2022). Hence, higher MD does not necessarily indicate immaturity or underdevelopment because face and fear biases are typical behavioural phenotypes at 8-months of age (Kataja et al., 2022; Leppänen, 2016). Rather, other brain structures, such as the amygdala (Tuulari et al., 2020) may facilitate this sustained AB to fear, despite the apparently less coherent WM microstructure. The link between infant behaviours and overall neurodevelopment may also explain the lack of findings in the ROI-based analyses.

Research suggests that WM development is asynchronous, region dependent (Barnea-Goraly et al., 2005; Warrington et al., 2022) and nonlinear up to 5 years of age (Stephens et al., 2020). Association fibres are the last to develop (Buyanova & Arsalidou, 2021; Huang et al., 2006) which could explain why the ROI-based analyses did not yield any results since the CNG, UF, and ILF are all association tracts and are still immature during the time of scanning at birth (Liu et al., 2022). Likewise, the pathways associated with DP FE were not found in the occipital lobe, but rather in the posterior CC, the external and internal capsule, SLF, and SFOF, which are sensorimotor and cognition pathways. This is in line with developmental patterns that show myelination occurs firstly in limbic fibres and develops anterior to posterior in commissural and projection fibres (Huang et al., 2006; Liu et al., 2022). Based on WM developmental patterns, we speculate that the heightened attention to fearful stimuli is maintained until the overall WM tracts are mature enough to facilitate semi-independent motoric exploration away from the caregiver (Callaghan et al., 2019; Peltola et al., 2013). Variance in neonatal WM microstructure may reflect individual differences in growth that is predictive of AB development later in infancy. However, further longitudinal research is needed to confirm the brain pathways related AB development throughout infancy.

### Limitations

This study adds to the understanding of infant development showing that WM microstructure at birth is predictive of infant attentional behaviour in infancy. The separate measuring time points of this study enhance understanding of the trajectories between neonatal WM microstructure and later behavioural phenotypes. However, the differing time points also complicate result interpretation. MRI measures at 8-months would have provided further insight into concurrent relationships between the infant brain and behaviour at this age, as well as the developmental changes since birth. Additionally, the limited sample size limits statistical power, especially within the sex-stratified analyses. Furthermore, the study cohort consists of a homogeneous Finnish, high SES population which may limit the findings generalizability to non-European or lower SES populations.

## Conclusions

Overall, the current study shows that neonatal brain structure is associated with behavioural development in the first year of life. More longitudinal studies with multiple follow-ups during infancy and early childhood are needed to confirm the hypotheses on general maturation of WM and later attentional biases to fear and how they deviate in pathology.

## Data Availability

Current EU and national legislation on personal data protection of sensitive data and the informed consents given by the study subjects do not permit open data sharing of imaging data or the derived measures. Investigators interested in research collaboration and obtaining access to the data are encouraged to contact FinnBrain board. Contact information of the board members or FinnBrain’s Principal Investigators are listed on the project website: https://sites.utu.fi/finnbrain/en/contact/.

## Supporting information

All supplementary materials

## Acknowledgements

We would like to warmly thank all FinnBrain families that participated in the study. We would also like to thank the research team that had a supportive role in the current study: Satu Lehtola for her help in data collection, Maria Lavonius for her help in recruiting the participants, Jani Saunavaara for implementing the MRI sequences, Riitta Parkkola for reviewing the MR images for incidental findings, and Jukka Leppänen for his expertise on the eye tracking experiment.

