## Supplementary material for "Neonatal White Matter Microstructure Predicts Infant Attention Disengagement from Fearful Faces": All supplementary materials

### 1. ROI results

#### Regression models

Table 1.1. Unadjusted regression models with fear bias scores and WM tracts

Abbreviations: Cingulate gyrus (cgc) and parahippocampal parts of the cingulum (cgh); UF: uncinate fasciculus; ILF: inferior longitudinal fasciculus

| | Adjusted $R^2$ | F (1,74) | p |
|---|---|---|---|
| **Left cgc** | -0.008 | 0.37 | 0.54 |
| **Right cgc** | -0.008 | 0.20 | 0.66 |
| **Left cgh** | -0.010 | 0.27 | 0.61 |
| **Right cgh** | -0.009 | 0.35 | 0.56 |
| **Left UF** | -0.011 | 0.15 | 0.70 |
| **Right UF** | -0.013 | 0.07 | 0.79 |
| **Left ILF** | -0.007 | 0.50 | 0.48 |
| **Right ILF** | -0.009 | 0.34 | 0.60 |

After adjusting for infant age at MRI from conception, age at ET from birth, sex, maternal age, education and BMI. None of the adjusted models significantly predicted fear bias as detailed in Table 5.

Table 1.2. Adjusted ANOVA models between fear bias and WM tracts.

| Model | Adjusted $R^2$ | F (1,74) | p |
|---|---|---|---|
| **Left cgc** | -0.090 | 0.36 | 0.95 |
| **Right cgc** | -0.090 | 0.36 | 0.95 |
| **Left cgh** | -0.091 | 0.35 | 0.95 |
| **Right cgh** | -0.091 | 0.35 | 0.95 |
| **Left UF** | -0.064 | 0.54 | 0.84 |
| **Right UF** | -0.085 | 0.39 | 0.94 |
| **Left ILF** | -0.063 | 0.54 | 0.84 |
| **Right ILF** | -0.061 | 0.56 | 0.83 |

Table 1.3. Pearson's correlations of variable of interests

Abbreviations: FA: Fractional Anisotropy; cgc: cingulate gyrus part of cingulum; cgh: parahippocampal part of cingulum; UF: uncinate fasciculus; ILF: inferior longitudinal fasciculus; L: left; R: right.

| | | **1.** | **2.** | **3.** | **4.** | **5.** | **6.** | **7.** | **8.** | **9.** | **10.** | **11.** | **12.** | **13.** | **14.** | **15.** |
|---|---|---|---|---|---|---|---|---|---|---|---|---|---|---|---|---|
| 1. Gestational age | | 1 | | | | | | | | | | | | | | |
| 2. Age at MRI | *r* | .495$^{***}$ | 1 | | | | | | | | | | | | | |
| | *p* | <.001 | | | | | | | | | | | | | | |
| 3. Age at ET | *r* | -.775$^{**}$ | -.338$^{**}$ | 1 | | | | | | | | | | | | |
| | *p* | <.001 | .003 | | | | | | | | | | | | | |
| 4. Maternal age | *r* | -.079 | -.051 | -.012 | 1 | | | | | | | | | | | |
| | *p* | .499 | .660 | .917 | | | | | | | | | | | | |
| 5. Maternal Education | *r* | -.105 | .058 | .047 | .192 | 1 | | | | | | | | | | |
| | *p* | .375 | .628 | .695 | .103 | | | | | | | | | | | |
| 6. Maternal BMI | *r* | -.100 | -.165 | -.021 | .260$^{*}$ | -.101 | 1 | | | | | | | | | |
| | *p* | .392 | .158 | .861 | .024 | .397 | | | | | | | | | | |
| 7. Face bias | *r* | -.020 | .100 | -.192 | -.002 | -.040 | .195 | 1 | | | | | | | | |
| | *p* | .864 | .389 | .099 | .988 | .735 | .094 | | | | | | | | | |
| 8. Fear bias | *r* | .057 | .021 | .001 | -.142 | -.131 | .104 | -.060 | 1 | | | | | | | |
| | *p* | .628 | .856 | .991 | .223 | .269 | .373 | .607 | | | | | | | | |
| 9. FA_cgc_L | *r* | .042 | .292$^{*}$ | .090 | -.005 | -.109 | -.084 | -.058 | .071 | 1 | | | | | | |
| | *p* | .719 | .010 | .443 | .969 | .358 | .474 | .618 | .544 | | | | | | | |
| 10. FA_cgc_R | *r* | .076 | .366$^{***}$ | -.050 | .015 | -.039 | -.020 | .098 | .052 | .572$^{***}$ | 1 | | | | | |
| | *p* | .513 | .001 | .671 | .898 | .744 | .868 | .398 | .656 | <.001 | | | | | | |
| 11. FA_cgh_L | *r* | -.013 | .359$^{***}$ | .034 | -.033 | .043 | .009 | .022 | .060 | .267$^{*}$ | .404$^{***}$ | 1 | | | | |
| | *p* | .909 | .001 | .774 | .776 | .719 | .939 | .848 | .607 | .020 | <.001 | | | | | |

| | | | 1. | 2. | 3. | 4. | 5. | 6. | 7. | 8. | 9. | 10. | 11. | 12. | 13. | 14. | 15. |
|---|---|---|---|---|---|---|---|---|---|---|---|---|---|---|---|---|---|
| 12. | FA_cgh_R | *r* | .033 | .295** | .040 | -.108 | -.148 | -.126 | .038 | .068 | .321** | .288* | .628*** | 1 | | | |
| | | *p* | .778 | .010 | .735 | .351 | .210 | .282 | .743 | .557 | .005 | .012 | <.001 | | | | |
| 13. | FA_UF_L | *r* | .286* | .661*** | -.157 | -.091 | -.155 | -.057 | -.046 | -.045 | .460*** | .464*** | .503*** | .400*** | 1 | | |
| | | *p* | .012 | <.001 | .179 | .434 | .189 | .625 | .694 | .700 | <.001 | <.001 | <.001 | <.001 | | | |
| 14. | FA_UF_R | *r* | .147 | .597*** | -.111 | .042 | .010 | .068 | .067 | -.031 | .321** | .469*** | .548*** | .368*** | .774*** | 1 | |
| | | *p* | .204 | <.001 | .345 | .716 | .930 | .564 | .567 | .788 | .005 | <.001 | <.001 | .001 | <.001 | | |
| 15. | FA_ILF_L | *r* | .049 | .465*** | .017 | -.094 | -.081 | -.009 | .098 | -.082 | .466*** | .455*** | .556*** | .352** | .655*** | .597*** | 1 |
| | | *p* | .672 | <.001 | .882 | .417 | .496 | .940 | .400 | .484 | <.001 | <.001 | <.001 | .002 | <.001 | <.001 | |
| 16. | FA_ILF_R | *r* | .292* | .587*** | -.077 | -.212 | -.019 | -.273* | -.037 | -.068 | .369*** | .389*** | .426*** | .513*** | .621*** | .593*** | .578*** |
| | | *p* | .011 | <.001 | .512 | .066 | .873 | .018 | .751 | .559 | .001 | <.001 | <.001 | <.001 | <.001 | <.001 | <.001 |

* $p \leq 0.05$, ** $p \leq .01$, *** $p \leq .001$ all significance levels are 2-tailed.

#### 2. Sex-stratified results

Sex-based stratification showed that males drove the overall sample results for both fear and happy conditions. In males, slightly higher MD values were found in superior and anterior fibres, towards the prefrontal cortex for DP fear (Figure S2.1, $p \leq .05$). Meanwhile, lower DP happy in males were still significantly associated with higher MD in the CC and possible corticospinal tract areas of the brain (Figure S2.2, $p \leq .05$). MD in males also showed a negative association with DP from neutral faces widespread across the brain (Figure S2.3, $p \leq .05$) including in the CC, bilateral external and internal capsules, bilateral SFOF, bilateral UF, left SLF, left posterior thalamic radiation. MD in males showed a positive association with face bias in the right external capsule (Figure S2.4, $p \leq .05$). No significant associations between any WM DTI metrics and attention measures were found in the female-only stratified sample ($p > .05$).

Previous findings regarding sex differences in early infancy brain development are mixed. One study found no sex-differences in infancy WM development, until later where females showed accelerated WM maturation at 5 years of age compared to males (Kumpulainen et al., 2023). Another study found that after controlling for total brain volume, neonatal males had larger total WM volume, whilst females had larger CC compared to males (Khan et al., 2024). However, in a longitudinal study of 6-, 12-, and 24-months old infants, males and females had the same CC area and thickness at 6 months, but males had a higher rate of CC growth from 6-24 months (Schmied et al., 2020). Hence, the current results provide further insight into the possibility that smaller CC at birth and slower CC development in males before 6 months may contribute to how attentional bias behaviour develops in male infants.

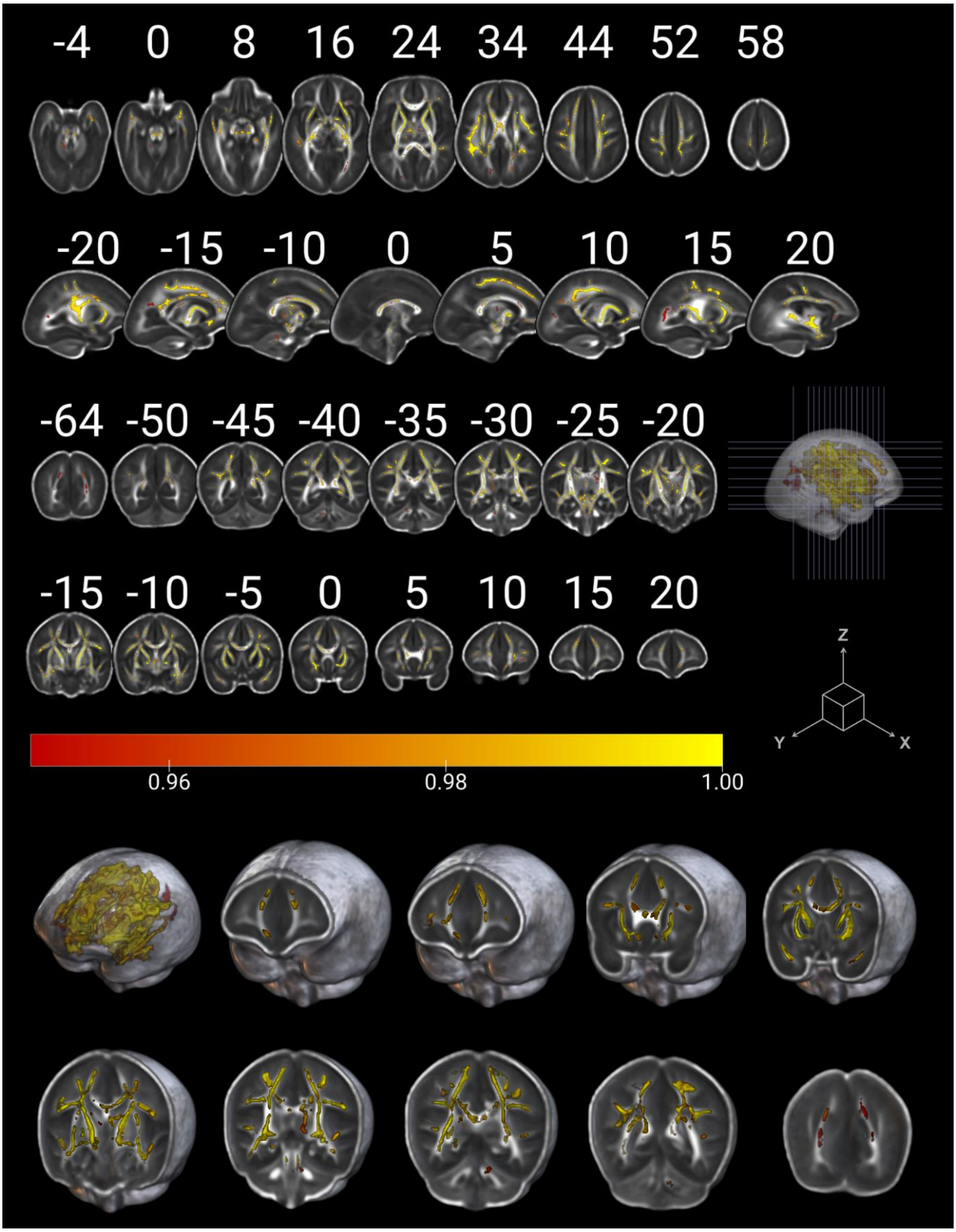


**Figure 2.1.** Higher mean diffusivity in male newborn brains is associated with lower attention disengagement from fearful faces at 8 months (threshold-free cluster enhancement (TFCE) corrected $p \leq 0.05$; across 5000 permutations). The colour bar represents $1 - p$.

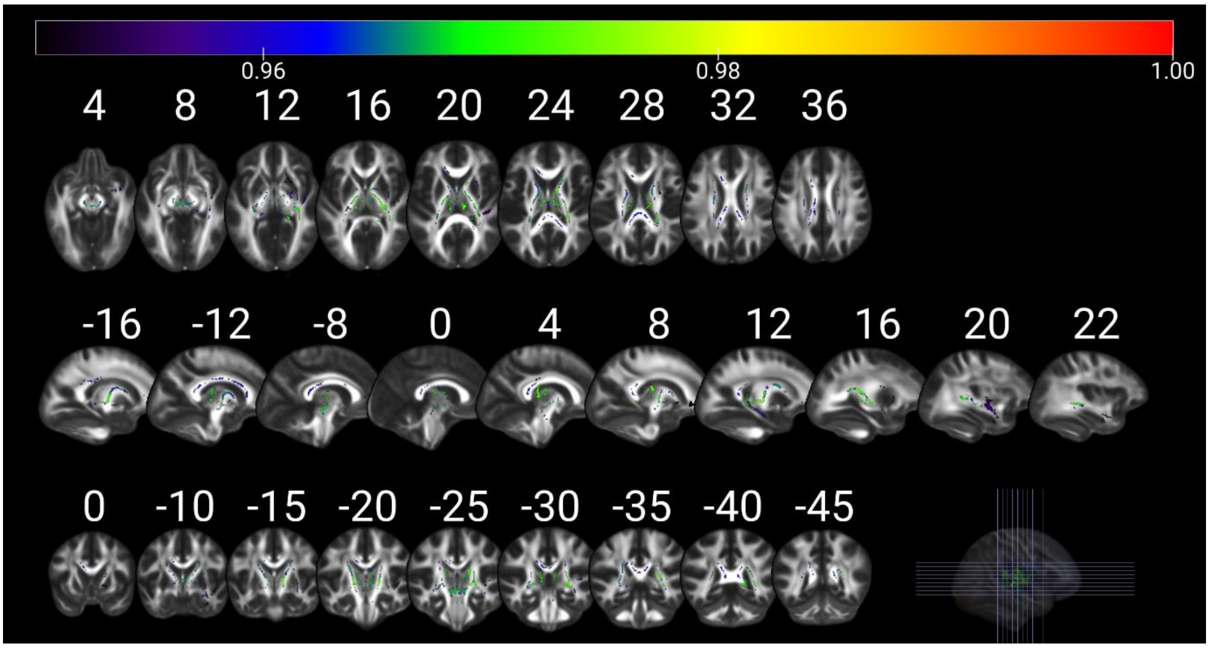


**Figure 2.2.** Higher mean diffusivity in male newborn brains is associated with lower attention disengagement from happy faces at 8 months ($p \leq 0.05$; 5000 permutations, multiple comparison corrections using threshold-free cluster enhancement). The colour bar represents $1 - p$.

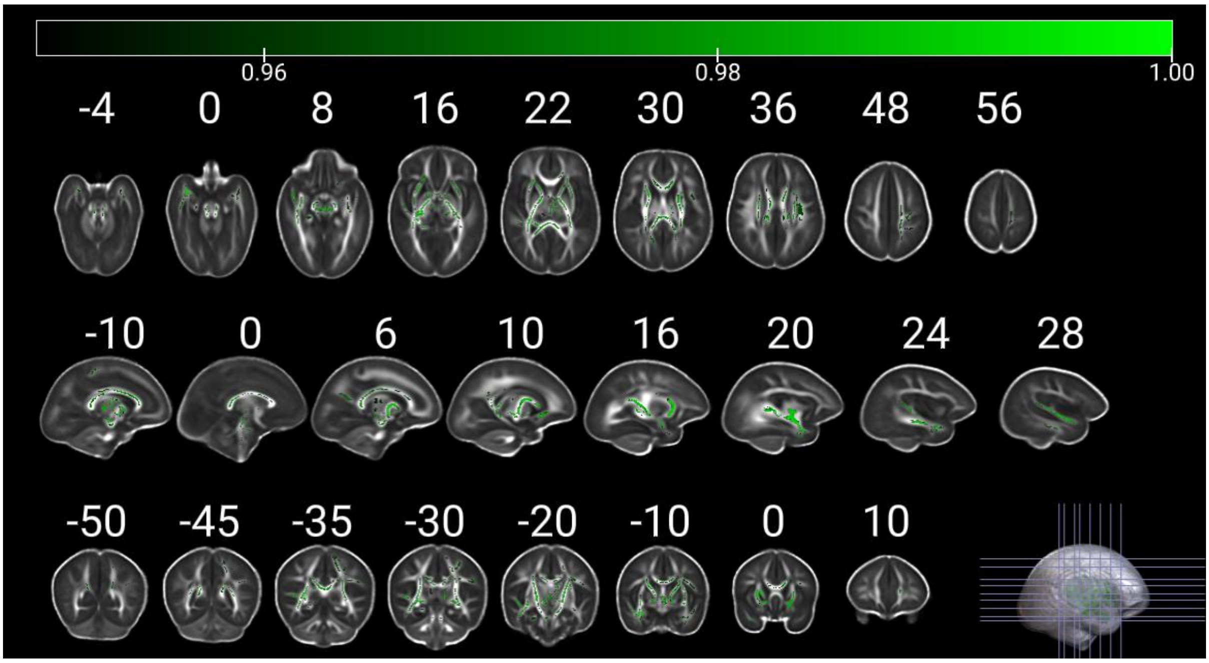


**Figure 2.3.** Higher mean diffusivity in male newborn brains is associated with lower attention disengagement from neutral faces at 8 months (threshold-free cluster enhancement (TFCE) corrected p ≤ 0.05; across 5000 permutations). The colour bar represents $1 - p$ .

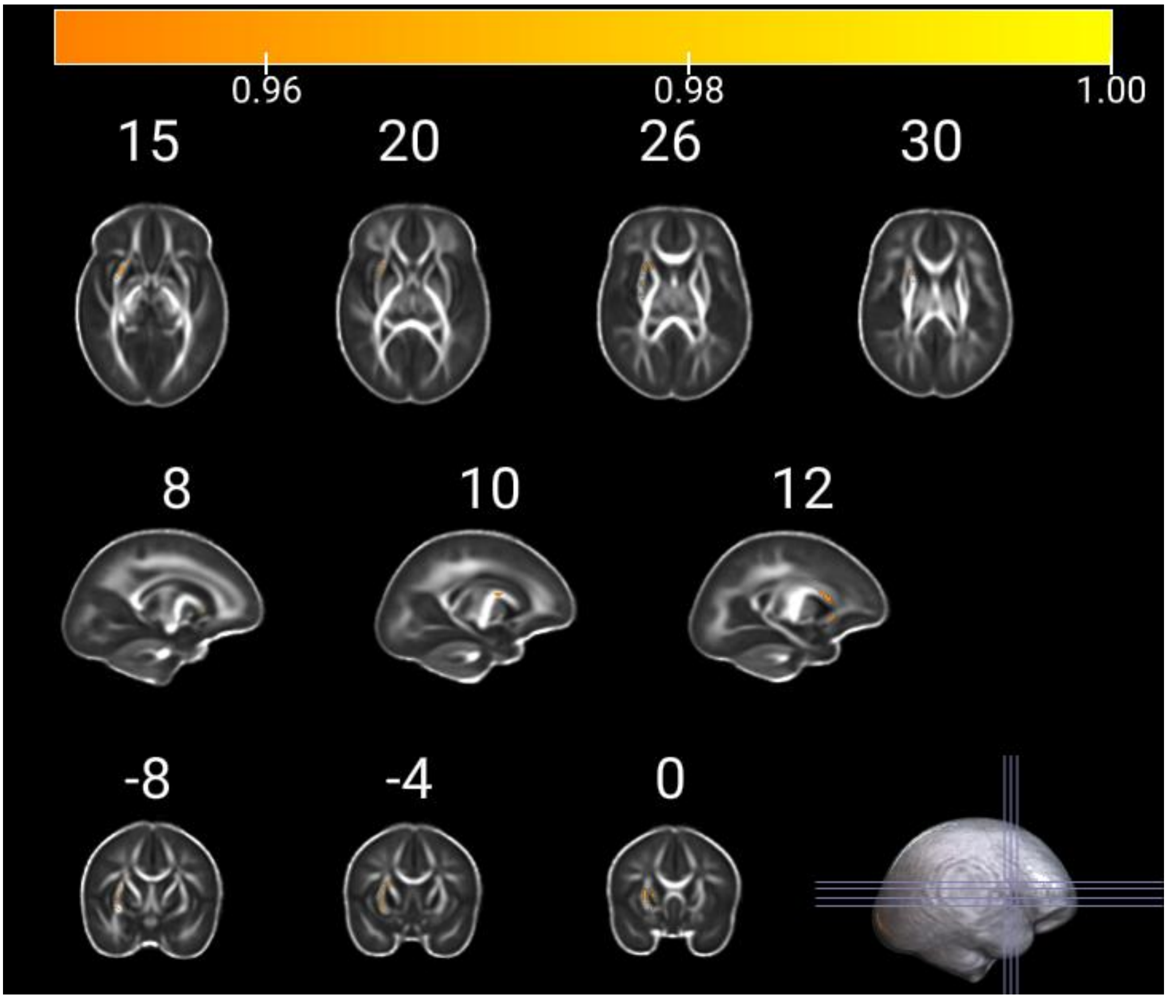


**Figure 2.4.** Higher mean diffusivity in male newborn brains is associated with higher face bias scores at 8 months (threshold-free cluster enhancement (TFCE) corrected p ≤ 0.05; across 5000 permutations). The colour bar represents $\mathbf{1 - \boldsymbol{p}}$.

### 3. Whole brain Voxelwise Sensitivity Analysis

#### DP FEAR

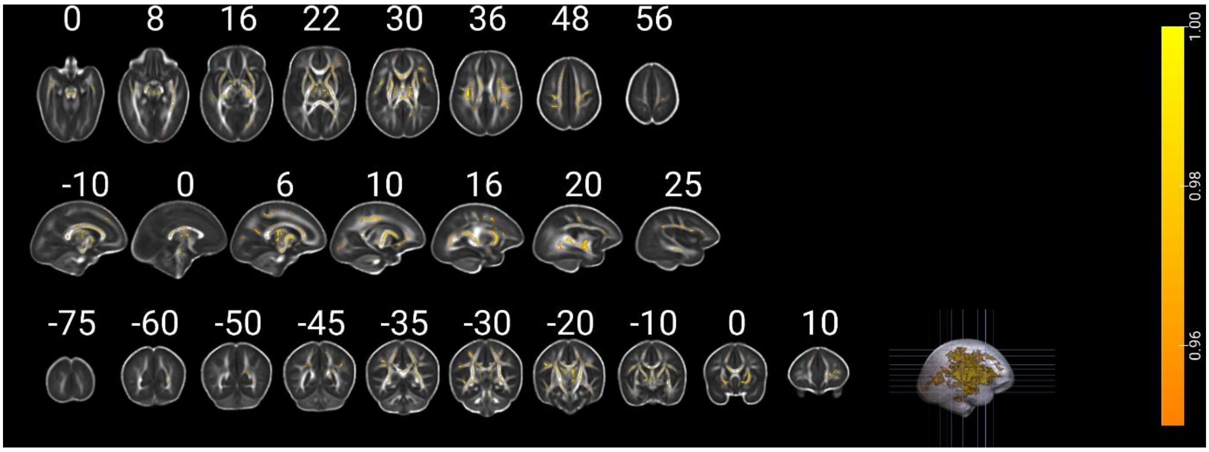


**Figure 3.1.** Higher mean diffusivity in newborn brains further controlling for birthweight is associated with lower attention disengagement from fearful faces at 8 months ($p \leq$ 0.05; 5000 permutations, multiple comparison corrections using threshold-free cluster enhancement). The colour bar represents 1−$p$.

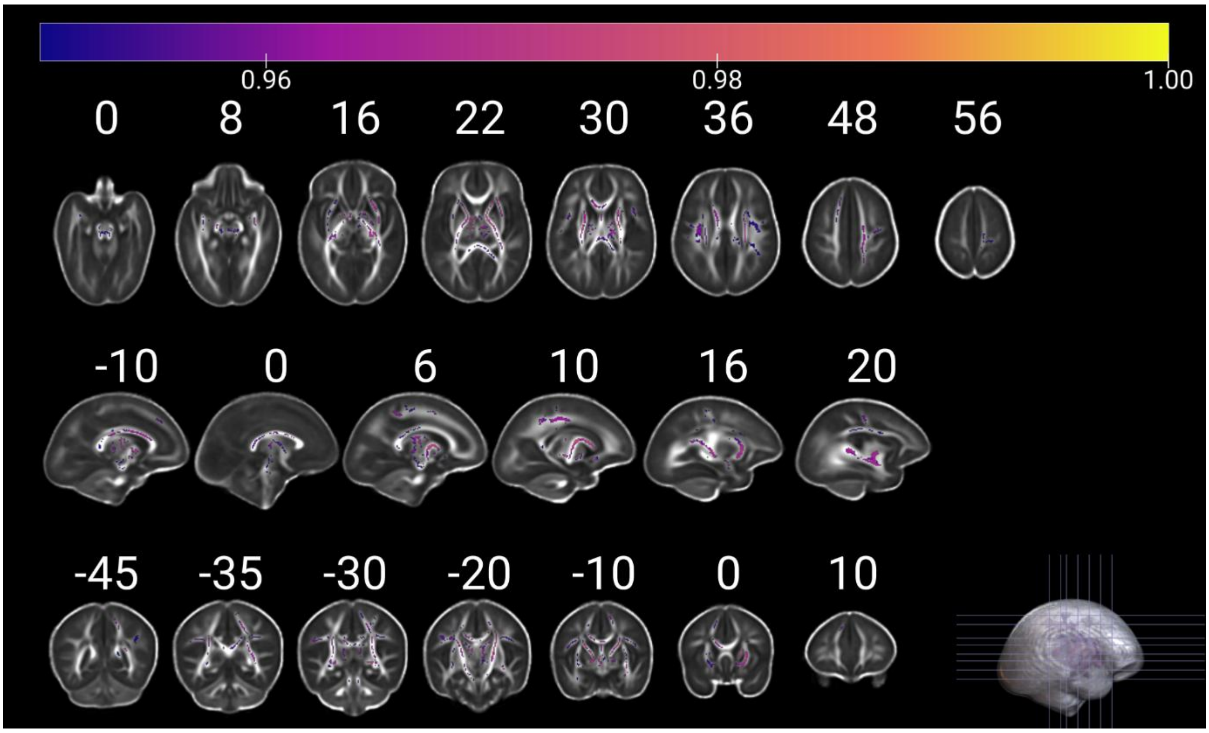


**Figure 3.2.** Higher mean diffusivity in newborn brains further controlling for infant head circumference is associated with lower attention disengagement from fearful faces at 8 months (threshold-free cluster enhancement (TFCE) corrected p ≤ 0.05; across 5000 permutations). The colour bar represents 1−$p$.

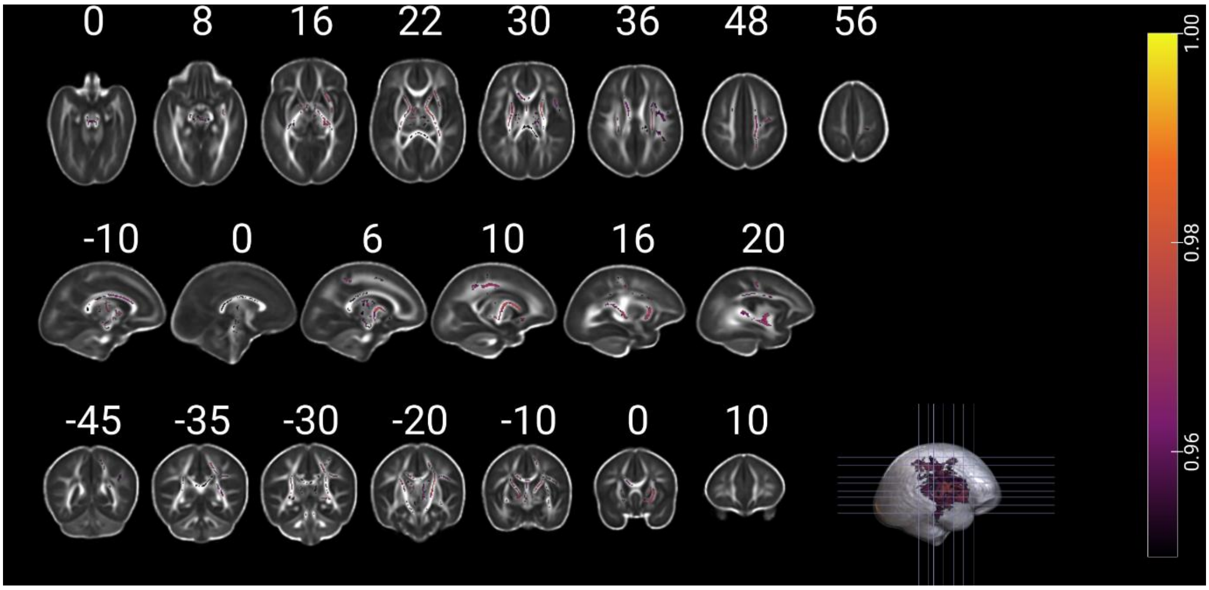


**Figure 3.3.** Higher mean diffusivity in newborn brains further controlling for alcohol is associated with lower attention disengagement from fearful faces at 8 months (threshold-free cluster enhancement (TFCE) corrected $p \leq 0.05$; across 5000 permutations). The colour bar represents 1–$p$.

DP Happy

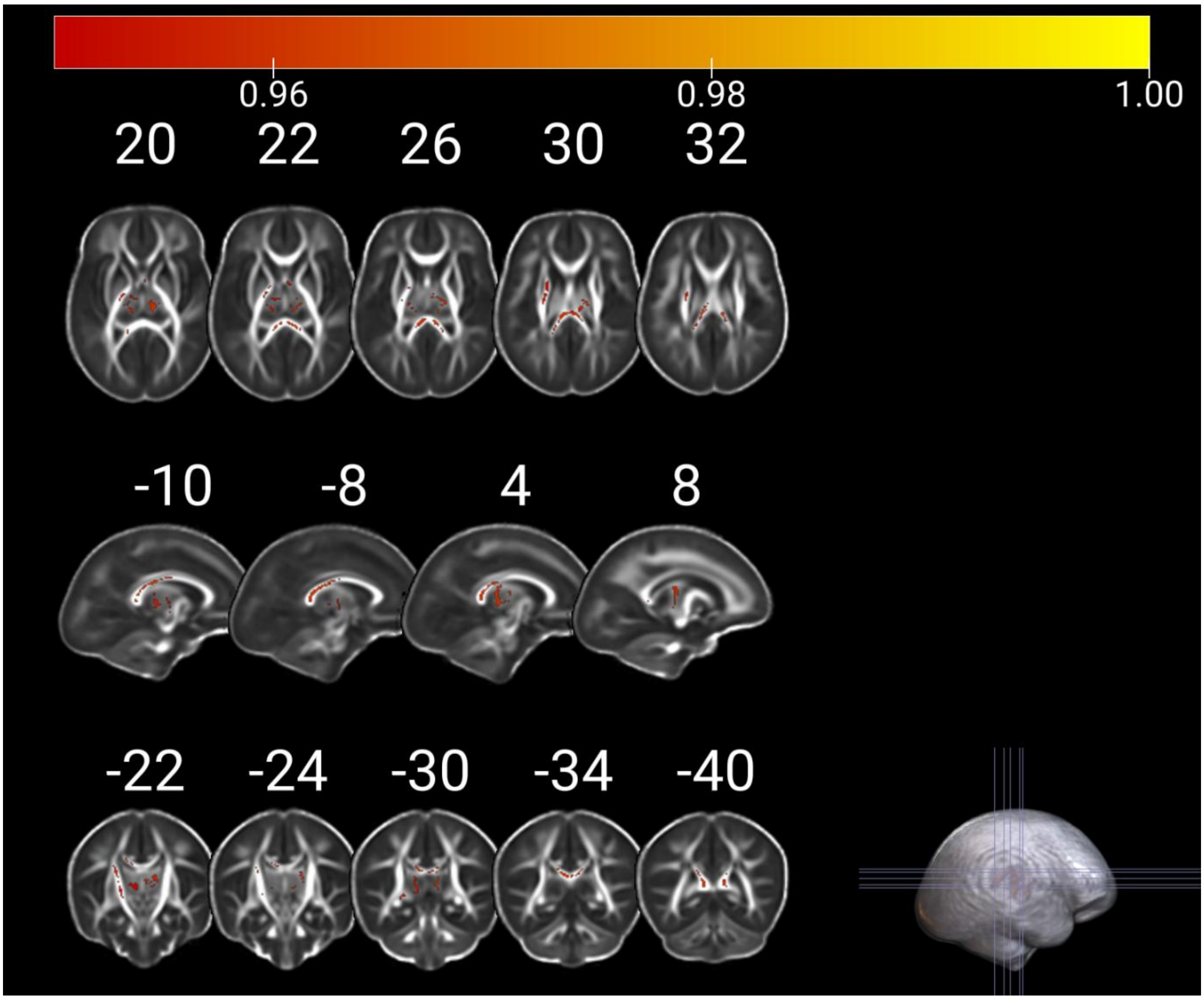


**Figure 3.4.** Higher mean diffusivity in newborn brains further controlling for birthweight is associated with lower attention disengagement from happy faces at 8 months (threshold-free cluster enhancement (TFCE) corrected p ≤ 0.05; across 5000 permutations). The colour bar represents 1–*p*.

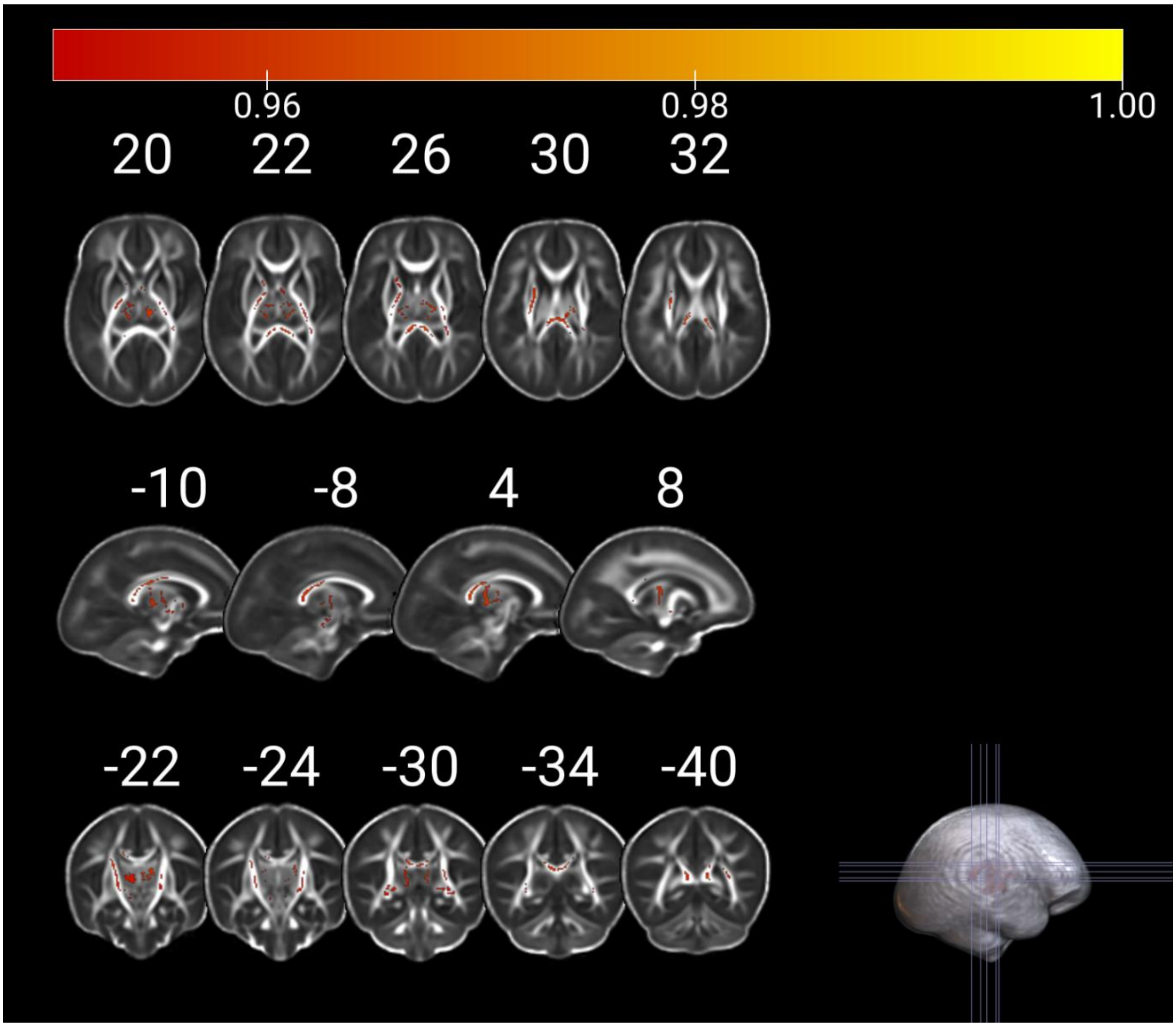


**Figure 3.5.** Higher mean diffusivity in newborn brains further controlling for maternal age is associated with lower attention disengagement from happy faces at 8 months (threshold-free cluster enhancement (TFCE) corrected $p \leq 0.05$; across 5000 permutations). The colour bar represents $1-p$.

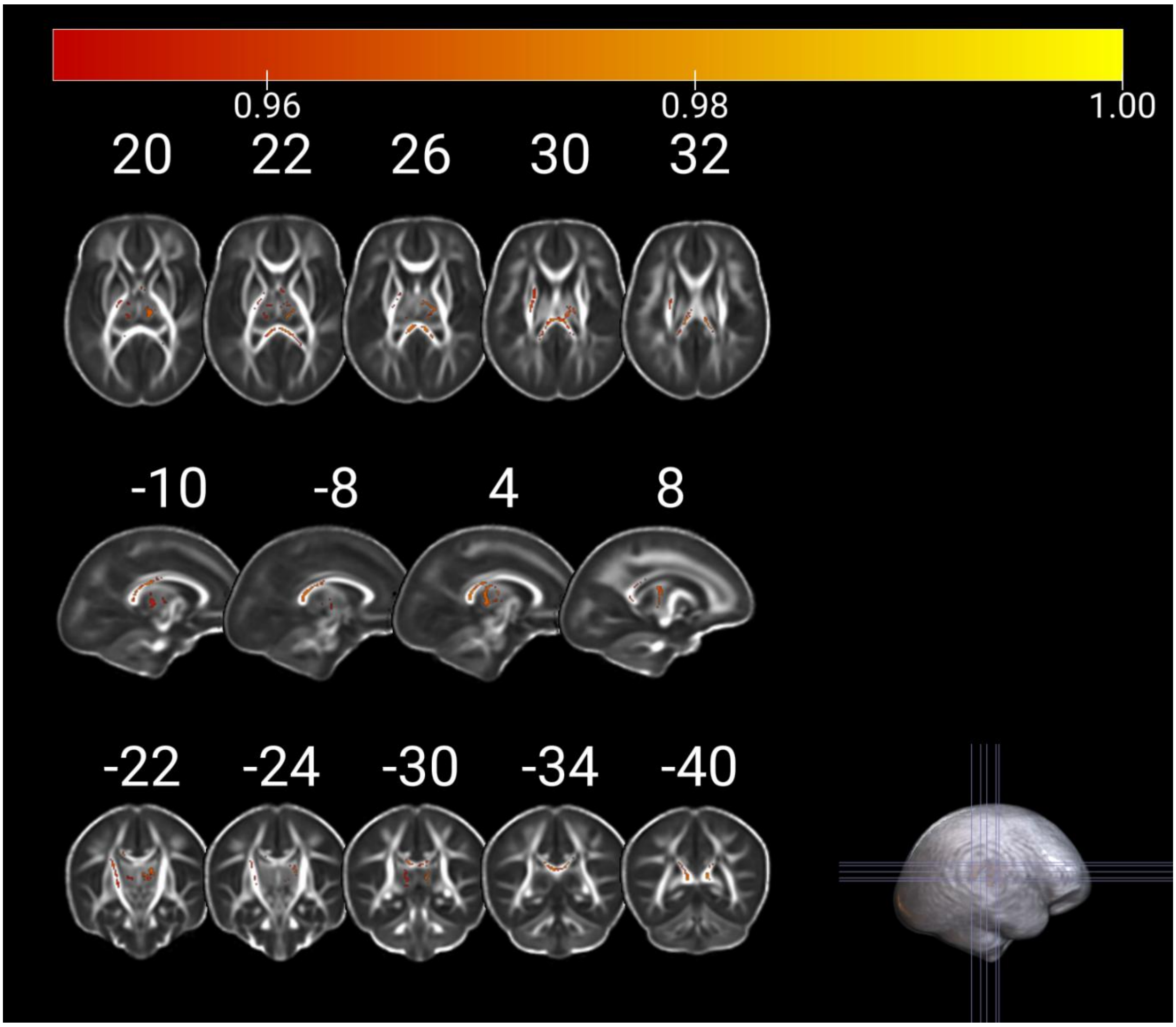

**Figure 3.6.** Higher mean diffusivity in newborn brains further controlling for alcohol is associated with lower attention disengagement from happy faces at 8 months (threshold-free cluster enhancement (TFCE) corrected $p \leq 0.05$; across 5000 permutations). The colour bar represents $1–p$.
